# Microbiomes of the bloom-forming alga, *Phaeocystis globosa*, are stable, consistently recruited communities with symbiotic and opportunistic modes

**DOI:** 10.1101/2022.02.02.478862

**Authors:** Margaret Mars Brisbin, Satoshi Mitarai, Mak A. Saito, Harriet Alexander

**Affiliations:** Biology Department, Woods Hole Oceanographic Institution; Marine Chemistry and Geochemistry Department, Woods Hole Oceanographic Institution; Marine Biophysics Unit, Okinawa Institute of Science and Technology

## Abstract

*Phaeocystis* is a cosmopolitan, bloom-forming phytoplankton genus that contributes significantly to global carbon and sulfur cycles. During blooms, *Phaeocystis* species produce large carbon-rich colonies, thus creating a unique interface for bacterial interactions. While bacteria are known to interact with phytoplankton—*e.g*., they promote growth by producing phytohormones and vitamins—such interactions have not been shown for *Phaeocystis*. Therefore, we investigated the composition and function of *P. globosa* microbiomes. Specifically, we tested whether microbiome compositions are consistent across individual colonies from four *P. globosa* strains, whether similar microbiomes are re-recruited after antibiotic treatment, and how microbiomes affect *P. globosa* growth under limiting conditions. Results illuminated a core colonial *P. globosa* microbiome—including bacteria from the orders Alteromonadales, Burkholderiales, and Rhizobiales—that was re-recruited after microbiome disruption. Consistent microbiome composition and recruitment is indicative that *P. globosa* microbiomes are stable-state systems undergoing deterministic community assembly and suggests there are specific, beneficial interactions between *Phaeocystis* and bacteria. Growth experiments with axenic and nonaxenic cultures demonstrated that microbiomes allowed continued growth when B-vitamins were withheld, but that microbiomes accelerated culture collapse when nitrogen was withheld. In sum, this study reveals interactions between *Phaeocystis* colonies and microbiome bacteria that could influence large-scale phytoplankton bloom dynamics and biogeochemical cycles.

## Introduction

Half of the photosynthesis on Earth is performed by phytoplankton, making them critical components of the global carbon cycle [1]. Like most organisms, phytoplankton do not act in isolation but instead interact with bacteria living in close proximity—either on phytoplankton cell surfaces or in the chemically influenced area surrounding phytoplankton (*i.e*., the phycosphere), a region analogous to the rhizosphere in plants [2]. While the phycosphere is not as extensively studied as the rhizosphere [3], it clearly hosts specialized, specific interactions with larger implications for marine ecosystem functioning and biogeochemical cycles [4]. For example, phycosphere bacteria can stimulate the growth of specific phytoplankton by producing growth hormones [5] or supplying a limiting nutrient, such as a B-vitamin [6–8] or fixed-nitrogen [9]. Conversely, phycosphere bacteria can manipulate phytoplankton communities by secreting algicidal molecules (*e.g*., roseobacticides) that selectively kill certain phytoplankton [10] or by competing for essential nutrients [11, 12]. Even small shifts in phytoplankton communities can alter how much organic carbon is recycled in the surface ocean or exported to deep water [13] since phytoplankton taxa vary in their C:N:P ratio [14], morphology, and sinking rates [15]. Therefore, phytoplankton-bacteria interactions are central to fully understanding phytoplankton dynamics and carbon cycling in the oceans. However, the stability, specificity, and currencies of exchange between phycosphere bacteria and diverse phytoplankton taxa are just beginning to be characterized.

The majority of our current knowledge regarding phytoplankton-bacteria interactions centers on interactions between diatoms and bacteria (reviewed in [16]) due to the ecological significance of diatoms on a global scale [17], as well as their experimental tractability (*e.g*., [5, 18]). Such studies have identified common bacterial inhabitants of phycospheres—including bacteria belonging to the orders Rhodobacterales, Flavobacteriales, Alteromonadales, and Oceanospirillales [18, 19]. Moreover, experiments with diatoms have demonstrated that phycosphere community assembly processes are mediated by both the phytoplankton and colonizing bacteria [20] and lead to highly reproducible bacterial community compositions [19]. However, it is unclear whether these traits extend to other phytoplankton taxa, particularly those that may be in competition with diatoms.

The Haptophyte genus, *Phaeocystis*, is an ideal system for extending studies of phytoplankton-bacterial interactions. *Phaeocystis* is a bloom-forming genus found from pole to pole. Notably, different bloom-forming species of *Phaeocystis* are specialized within latitudinal or thermal niches, often competing with diatoms [21]. *P. globosa* competes with diatoms to be the dominant spring bloom constituent in the North Sea, where it is considered a nuisance alga [22]. *P. globosa* is further distributed across temperate, subtropical, and tropical regions globally and has more recently started causing harmful algal blooms in Southeast Asia [23, 24]. The two polar species, *P. pouchetii* and *P. antarctica*, contribute to spring blooms in the Arctic / N. Atlantic oceans and the Southern Ocean, respectively [21]. In the N. Atlantic, *P. pouchetii* has begun to regularly replace diatoms in the spring bloom [25] and in the Southern Ocean, there is continued interannual variability as to whether *P. antarctica* or diatoms dominate spring blooms [26]. While not yet systematically studied, existing evidence supports the hypothesis that bacterial interactions could influence *Phaeocystis* blooms: consistent bacterial communities have been found in association with discrete samples of two *P. globosa* strains studied independently [27, 28] and *P. antarctica* blooms have been shown to strongly influence bacterioplankton community composition [29].

Bloom-forming *Phaeocystis* species share the ability to form large colonial structures (up to a centimeter in diameter) that are made up of thousands of cells embedded in a mucopolysaccharide matrix [30]. The carbon-rich colony substrate creates a unique physical and chemical interface for interaction between bacteria and *Phaeocystis* [31]. Bacteria are attracted to polysaccharides produced by phytoplankton [32] and the specific polysaccharides excreted by a phytoplankton cell contribute to which bacteria it encounters or is colonized by [18, 33, 34]. The concentrated availability of distinct polysaccharides in the *Phaeocystis* colony matrix, along with other chemical properties—such as the presence of amino sugars [35] and abundant dimethylsulfoniopropionate (DMSP) [36]—suggests a unique bacterial consortium would be attracted to and thrive in the colonial *Phaeocystis* microenvironment [18]. However, given the extensive functional redundancy among bacteria [37], it remains to be determined how specialized bacterial communities associated with *Phaeocystis* colonies are relative to other plankton and whether *Phaeocystis* microbiomes are stable-state systems.

In this study, we tested the specificity and stability of bacterial communities associated with *Phaeocystis* colonies by evaluating the microbiomes of individual colonies from replicate culture batches of four *P. globosa* strains that were isolated from globally wide-ranging locations across several decades. We further tested the reproducibility of *P. globosa* colony microbiome compositions by removing original microbiomes with antibiotics and evaluating compositions of recruited microbiomes after inoculation with bacterial communities in seawater. Finally, we tested two possible functional roles for *P. globosa* microbiomes by challenging axenic and nonaxenic cultures with nitrogen and vitamin stress. Nitrogen deprivation was chosen because nitrogen availability is a key factor believed to influence temperate and subtropical *Phaeocystis* blooms [25, 38] and because fixed-nitrogen is one currency of exchange between phycosphere bacteria and some diatoms [39]. B-vitamin limitation was chosen because (i) the availability of B-vitamins affects phytoplankton bloom dynamics [40, 41], (ii) *P. globosa* requires vitamins B_1_ and B_12_ to grow [42], and (iii) phycosphere bacteria can be sources of B-vitamins [7, 8, 43]. Overall, we found *P. globosa* colonies hosted remarkably consistent bacterial communities, both before antibiotic treatment (original microbiomes) and in the recruitment trial. These results demonstrate that, like many diatoms, *Phaeocystis* colonies regulate convergent bacterial community assembly towards a stable and reproducible core microbiome, albeit a different core microbiome than previously observed for other phytoplankton groups. Furthermore, the full consortia of co-cultured bacteria (in media and associated with colonies) differentially impact *P. globosa* growth depending on environmental conditions.

## Methods

### *Phaeocystis globosa* culture sources

We included four strains of *Phaeocystis globosa* in this study to investigate the consistency of *Phaeocystis-associated* bacterial community compositions across strains. Three strains—CCMP1528, CCMP629, and CCMP2710—were obtained from the National Center for Marine Algae and Microbiota (NCMA, Boothbay, ME, U.S.A.). We isolated the fourth strain in January 2019 from the Kuroshio current in the East China Sea (28.22 °N, 127.66 °E) by collecting surface water (5 m depth) and filtering it through 63 μm mesh-size nylon mesh before enriching with 10% L1-Si media [44]. Nutrient enrichment triggered a bloom of colonial *P. globosa* and individual *P. globosa* colonies were subsequently isolated, rinsed with sterile artificial seawater, and used to establish mono-algal cultures of a new *P. globosa* strain, referred to as “Kuroshio” herein. Together, these four strains represent wide geographic diversity and have been maintained in culture for widely varying times, from several decades to just a few years (Fig. 1).

**Figure 1.**
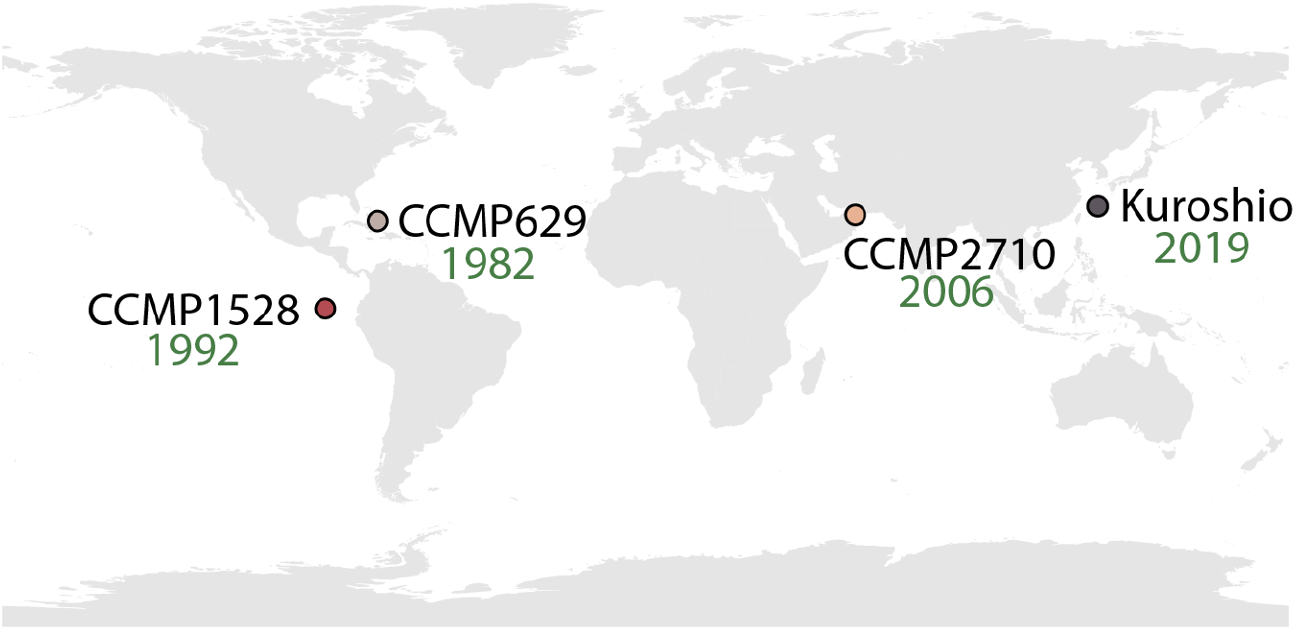
Geographic origins of the four strains of *Phaeocystis globosa* included in the study. The year that each strain was originally isolated is displayed below each strain’s name. “CCMP” strains were obtained from the National Center for Marine Algae and Microbiota in Boothbay, Maine, U.S.A. The Kuroshio strain was isolated for this study.

### Evaluation of *P. globosa* colony and media microbiomes

Cultures of each *P. globosa* strain were synchronized by inoculating 35 ml of L1-Si media in triplicate 50 ml plastic culture flasks with 1 ml of maintenance culture, incubating for five days, and transferring 1 ml to a newly prepared culture flask. Cultures were incubated in a TOMY digital biology CLE-series plant growth chamber with cool-white fluorescent lighting (16 h light / 8 h dark diurnal light cycle, ~ 50 μmol m^-2^s^-1^ light) maintained at 22°C. After five such transfers, individual colonies and free-living cells were harvested for DNA extraction. Three colonies from each culture replicate were rinsed three times with sterile artificial sweater (ASW) before careful transfer (with <10 μl of ASW) to an Axygen Maxymum Recovery PCR tube with 30 μl 10% mass:volume Chelex 100 resin beads in PCR-grade water. The colony and Chelex mixtures were vortexed for 10 s, incubated at 96°C for 20 minutes, vortexed again for 10 s, and placed on ice [45]. Free-living bacterial cells in *Phaeocystis* media were collected by gravity-filtering each culture replicate through 100-, 70-, and 40-μm mesh-size cell strainers to remove colonies and then filtering under gentle vacuum pressure through 3.0-μm (to remove free-living *Phaeocystis* cells) and 0.2-μm pore-size polytetrafluoroethylene (PTFE) filters (Millipore) to collect bacterial biomass. DNA was extracted from the 0.2-μm pore-size filters by following the manufacturer’s protocol for the DNeasy PowerSoil kit (Qiagen), including the optional heating step for enhanced cell lysis. The v3-v4 region of the 16S rRNA gene was amplified from extracted DNA from both extraction methods following the Illumina guide for “16S Metagenomic Sequencing Library Preparation” (https://support.illumina.com/content/dam/illumina-support/documents/documentation/chemistry_documentation/16s/16s-metagenomic-library-prep-guide-15044223-b.pdf). The 16S rRNA gene amplicon sequencing libraries were sequenced with 300×300-bp v3 paired-end chemistry on the Illumina MiSeq platform.

### Evaluation of recruited bacterial communities following antibiotic disruption of *Phaeocystis* microbiomes

To determine whether *Phaeocystis* strains reproducibly recruit specific bacteria to their microbiomes, the original microbiomes of each strain were disrupted and then each strain was exposed to bacterial assemblages in seawater before microbiome composition was assessed a second time. Original microbiomes were disrupted through antibiotic treatment: 1 ml of actively growing cultures were transferred to 50 ml culture flasks with 35 ml of L1-Si media and 1 ml of an antibiotic cocktail with 12 000 units/ml Penicillin G, 300 units/ml Polymyxin B, 50 μg/ml Chloramphenicol, 60 μg/ml Paromomycin and 6.25 mg/ml Cephalexin. After 48 h, 1 ml of each antibiotic-treated culture was transferred to L1-Si media without antibiotics and allowed to recover from antibiotic treatment for five five-day culture cycles. Microbiome disruption was checked by incubating aliquots of each antibiotic-treated culture with Hoechst 33342 DNA-binding dye and imaging each aliquot with confocal laser scanning fluorescence microscopy (Zeiss LSM 780). Additionally, aliquots of antibiotic-treated cultures were transferred to Difco Marine Broth 2216 (BD Biosciences) at the end of each ensuing culture cycle and monitored for bacterial growth by light microscopy (Olympus CKX53). No bacteria were observed in antibiotic-treated cultures with fluorescent microscopy, nor was bacterial growth detected with light microscopy in marine broth incubations. Henceforth, antibiotic-treated cultures were considered axenic.

Inoculating seawater for microbiome recruitment trials was collected outside the coral lagoon of Okinawa Island (Japan) by bucket cast and filtered through 63 μm-mesh-size nylon mesh before transport back to the lab. Inoculating seawater was further filtered through 3 μm- and 0.45 μm-pore-size PTFE filters to remove eukaryotic cells. Two 50 ml replicates of the inoculating seawater were filtered through 0.2 μm pore-size filters to assess the initial composition of bacterial communities inoculating *Phaeocystis* cultures. L1-Si media for recruitment trials was prepared with 50% inoculating seawater and 50% sterile artificial seawater. Culture replicates for microbiome recruitment trials were initiated with 1 ml of actively-growing axenic *Phaeocystis* cultures in triplicate flasks. The bacterial community was allowed to equilibrate over five consecutive five-day culture cycles (transferring every fifth day) before *Phaeocystis* colonies (five from each replicate) and free-living media-associated bacteria were harvested for DNA extraction and amplicon sequencing following the methods described above.

### Bioinformatics and statistical analyses

Sequencing for this project was completed using two full Illumina Miseq flow cells. Resulting sequencing reads were quality-filtered and Amplicon Sequence Variants (ASVs) were identified using the DADA2 denoising algorithm [46], implemented within the QIIME 2 framework (v2020.6, [47]), which also removes chimera and singleton sequences. Sequencing reads from each of the MiSeq runs were denoised independently with results merged after denoising, and ASVs clustered at 99% identity. Taxonomy was assigned to representative sequences with a naive Bayes classifier trained on the SILVA 99% 16S sequence database v138 [48] using the QIIME 2 feature-classifier plug-in [49]. The resulting ASV count table and taxonomic assignments were exported to the R statistical environment (v4.0.3, [50]) and further analyzed with the phyloseq (v1.34, [51]), CoDaSeq (v0.99.6, [52]), and vegan (v2.5-7, [53]) packages.

### Effects of microbiomes on *P. globosa* growth under limiting culture conditions

To assess impacts of microbiomes on *Phaeocystis* growth, we monitored axenic and nonaxenic *Phaeocystis* cultures under nutrient-replete and two limiting conditions: nitrogen limitation and B-vitamin limitation. L1-Si media was prepared without nitrate to induce nitrogen limitation and L1-Si media was prepared without the L1 vitamin solution, which contains thiamine (vit. B_1_), biotin (vit. B_7_), and cobalamin (vit. B_12_), to induce B-vitamin limitation [44]. Triplicate 50 ml sterile plastic culture flasks were prepared with 35 ml of each media type (L1-Si replete, L1-Si with no nitrate, and L1-Si with no B-vitamins) in a laminar flow hood sterilized with UV light and cleaned with ethanol. One ml of actively growing axenic or nonaxenic *P. globosa* was transferred to each flask so there were three replicates of axenic and nonaxenic cultures for each of the four strains (total of 72 flasks per culture cycle). Growth was monitored daily by removing 1 ml from each flask, homogenizing by pipette mixing, and measuring *in vivo* Chlorophyll *a* fluorescence with a Turner Designs Trilogy Fluorometer. Transfers to fresh media were completed for five culture cycles (5–7 days each, depending on growth) for each treatment, except for nonaxenic cultures in L1-Si media without nitrate, as growth ceased after the third culture cycle. All transfers and aliquots were made inside the clean laminar flow hood. Following each culture cycle, an additional 100 μl aliquot was taken from each flask, transferred to 1 ml sterile marine broth in a 12-well cell-culture plate, and monitored for bacterial growth for 72-hours. No bacterial growth was observed from aliquots taken from axenic cultures, whereas bacterial growth was always observed after 24-hours in aliquots from nonaxenic cultures, thus confirming axenic and non-axenic cultures, respectively.

## Results

### Sequence processing

Overall, amplicons from 100 samples (*n* = 34 original single-colony microbiomes, *n* = 12 original media microbiomes, *n* = 40 recruited single-colony microbiomes, *n* = 12 recruited media microbiomes, *n* = 2 inoculating seawater communities) were sequenced for this project, generating a total of 17 328 537 sequencing reads (median total sequences per sample = 147 351, Fig. S1). Raw sequencing data is available from the NCBI SRA under accession PRJN779092. Following quality-filtering, denoising, and chimera and singleton removal, 13 435 004 sequencing reads remained (median remaining sequences per sample = 121 035, Fig. S1). One concern when studying phytoplankton microbiomes is the proportion of sequencing resources allocated to non-target chloroplast 16S genes instead of target bacterial 16S genes. Lower input, single-colony samples yielded a larger proportion of chloroplast sequences than media or seawater samples: single colony samples had a mean of 22% bacterial sequences compared to 85% and 99.5% bacterial sequences for media and seawater samples, respectively (Fig. S2). However, following removal of chloroplast sequences, rarefaction curves for all samples reached ASV richness saturation (Fig. S3), indicating an adequate number of bacterial sequences remained for analysis due to sufficient sequencing depth. Chloroplast sequences were excluded from ensuing analyses.

### Diversity of original and recruited *P. globosa* microbiomes

The inoculating bacterial community for the recruitment trial was significantly more diverse than original and recruited *Phaeocystis* microbiomes (Fig. 2; pairwise Wilcox test, *p* < 0.05). In addition, original and recruited colony microbiomes were significantly less diverse than original and recruited media microbiomes (Fig. 2; pairwise Wilcox test, *p* < 0.001). In both these cases, colony microbiomes had about half the observed richness as media microbiomes.

**Figure 2.**
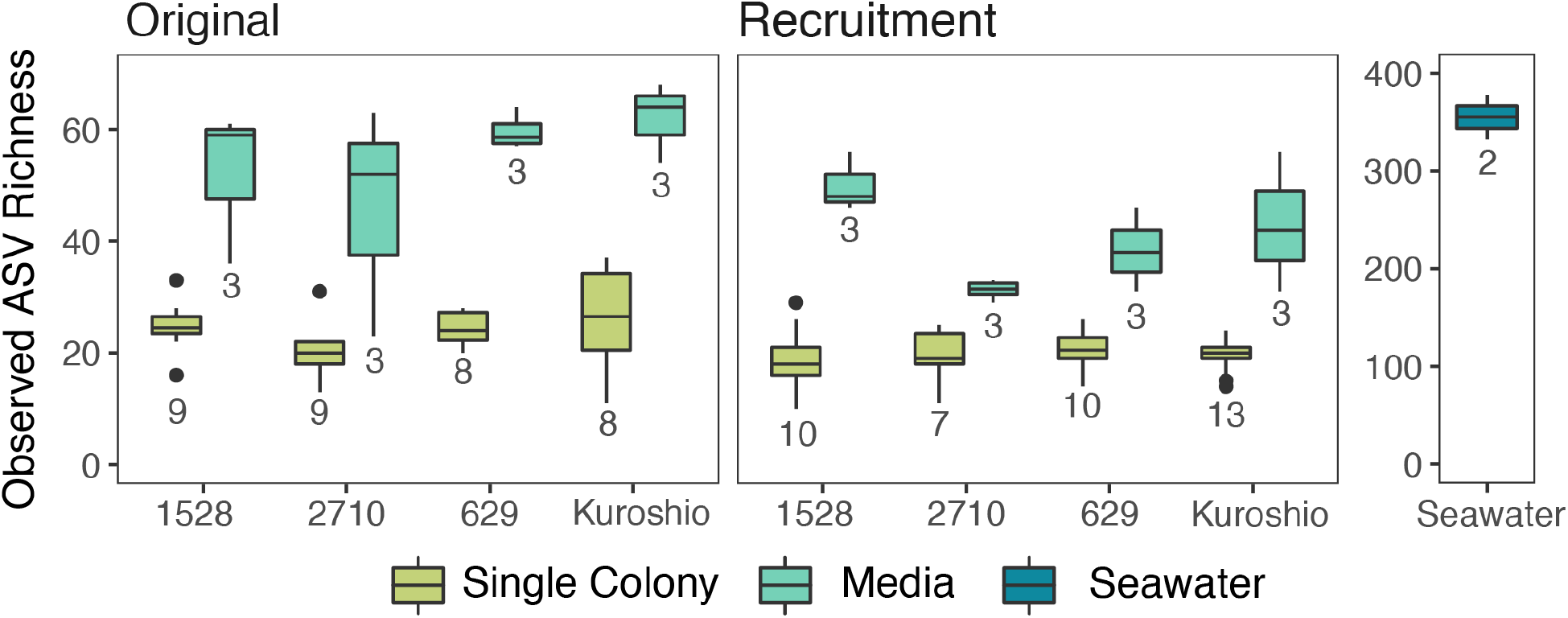
Observed richness of Amplicon Sequence Variants (ASVs) in original and recruited bacterial communities associated with individual *Phaeocystis* colonies, *Phaeocystis* culture media, and inoculating seawater used in recruitment trials. Results from the four *Phaeocystis globosa* strains are plotted separately along the x-axis. The number of biological replicates (*n*) for each sample type is included below the corresponding box. Note that the observed richness in the inoculating seawater is plotted on a separate y-axis due to the much higher diversity in these samples. Boxes denote the 25^th^ to 75^th^ percentile ranges for each sample type; the center bar is the median; whiskers show 1.5 times the interquartile range above the 75^th^ percentile and below the 25^th^ percentile; outlier points are values outside 1.5 times the interquartile range above the 75^th^ percentile and below the 25^th^ percentile. Media-associated bacterial communities are significantly more diverse (higher richness) than colony-associated bacterial communities (pairwise Wilcox test, *p* < 0.001).

The beta diversity between different sample types was computed by normalizing ASV sequence read counts to a centered log-mean ratio (CLR) using the CoDaSeq package and calculating Euclidean distances between community compositions in each sample with the phyloseq package. This combination of CLR transformation and Euclidean distance is known as the Aitchison distance and minimizes bias inherent to compositional data, such as amplicon data [52]. A Principal Coordinates Analysis (PCoA) performed on Aitchison distances revealed four clearly separated clusters: inoculating seawater communities, original media microbiomes, recruited media microbiomes, and colony-associated microbiomes (Fig. 3a). When sample points are colored by trial (original or recruitment), the colony-associated community cluster appears split by trial, but original and recruited colony microbiomes were not significantly different (Fig. 3a). Pairwise Permutational Multivariate Analysis of Variance (PERMANOVA) tests using the adonis2 function from the vegan package (999 permutations; [53]) showed original and recruited colony-associated communities were both significantly different from media and seawater-associated communities (*p* < 0.05) but not from each other, whereas recruited and original media-associated communities were both significantly different from the colony and seawater-associated communities as well as from each other (*p* < 0.05).

**Figure 3.**
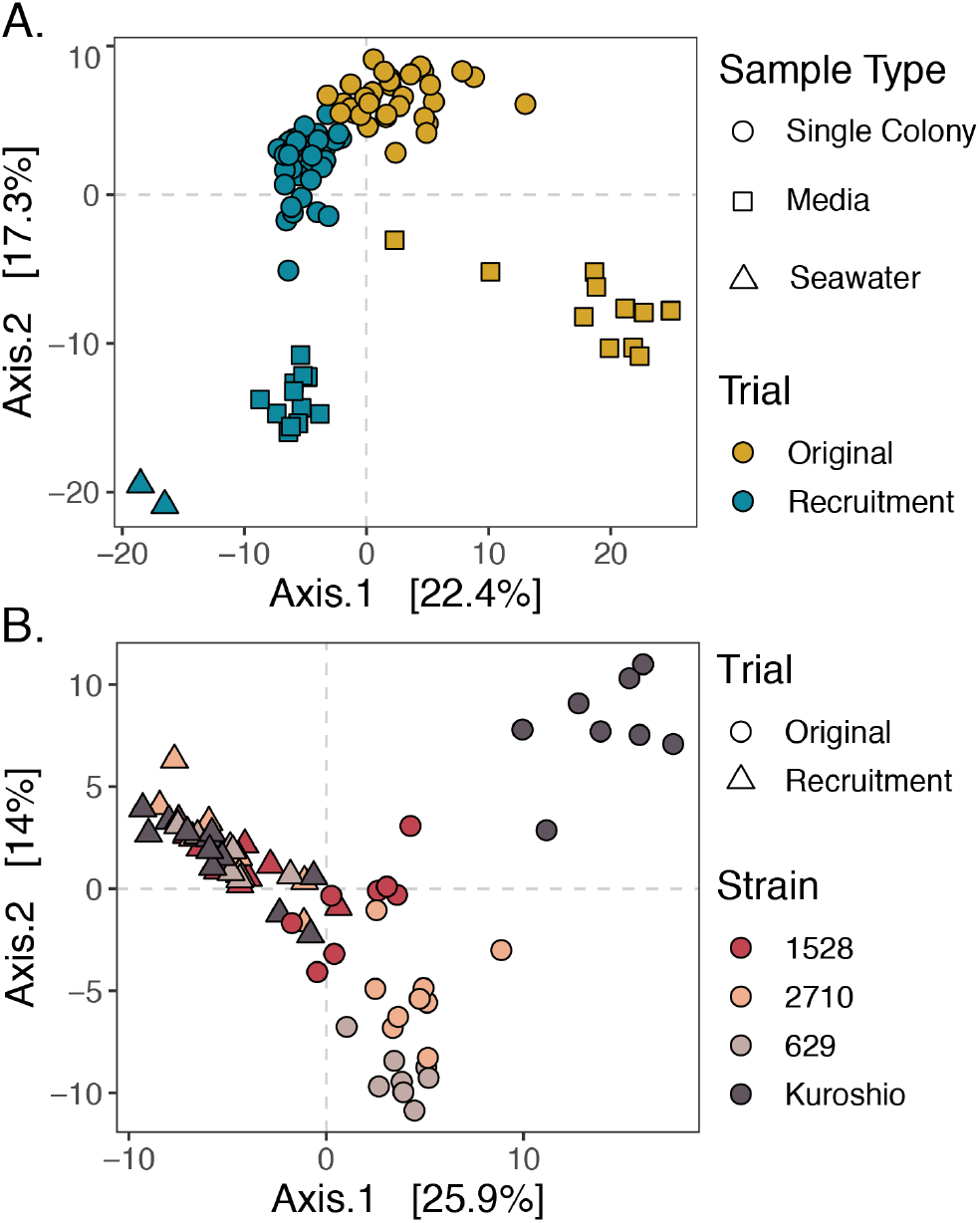
Principal Coordinates Analysis (PCoA) of Aitchison distances between bacterial communities associated with individual *Phaeocystis* colonies, *Phaeocystis* media, and inoculating seawater. (A) PCoA of Aitchison distance between all samples with point shape representing sample type (single colony, media, or seawater) and point color representing trial (original or recruitment). Four main clusters are visible: colony-associated communities, original media-associated communities, recruited media-associated communities, and communities in seawater. Coloration reveals that the colony-associated community cluster is split by trial. (B) PCoA of Aitchison distances between single-colony samples only, with point shape representing trial (original or recruitment) and color representing *P. globosa* strains. Samples separate by strain among original colony-associated communities, especially in regards to the Kuroshio colonies, but this differentiation collapses among recruited colony samples. *Phaeocystis* strain-level differences in colony microbiomes were significant among the original colony samples (*p* < 0.001, PERMANOVA, 999 permutations), but not among the recruitment trial colony samples (*p* = 0.117, PERMANOVA, 999 permutations).

To evaluate the consistency of colony microbiome compositions by host strain, data were subset to include only single-colony samples, and the Aitchison distances and PCoA were recomputed (Fig. 3b). This PCoA showed original colony microbiomes were differentiated by strain. Specifically, original colony microbiomes from the Kuroshio strain, which was isolated most recently and was not kept at a culture facility, cluster separately from the other original colony microbiomes. However, this separation by strain collapses when comparing microbiomes recruited to colonies after antibiotic treatment (Fig. 3b). Moreover, PERMANOVA tests by strain performed separately for original and recruited colony microbiomes showed strain-level differences were only significant (*p* < 0.001) among original microbiomes.

### Composition of original and recruited *P. globosa* microbiomes

To further examine ASVs composing *Phaeocystis* colony and media microbiomes, data were subset to exclude ASVs only present in seawater samples. Subsetting reduced the total unique ASV count from 781 to 341. When the relative abundance of each remaining ASV was plotted for each sample, trends in original and recruited colony- and media-associated microbiomes were readily apparent (Fig. 4). Original and recruited colony microbiomes were remarkably consistent across strains, with most colony microbiomes including the same ASVs from the orders Alteromonadales, Burkholderiales, and Rhizobiales. Original colony microbiomes also consistently included Caulobacterales ASVs, whereas recruited colony microbiomes consistently included an Oceanospirillales ASV that was not in original colony or media-associated microbiomes but was in recruited media microbiomes and in inoculating seawater. In addition, the plot highlighted two ASVs in the order Oceanospirillales contributing to the Kuroshio strain’s distinct original colony microbiomes (Fig. 3b). Original media-associated microbiomes were most consistent, with almost all ASVs detected being found in at least 9 out of 12 samples. Notably, recruited ASVs were not always also detected in seawater. This is likely due to recruited ASVs representing bacteria with very low abundance in the inoculating seawater, especially since this seawater was filtered to remove particles and eukaryotic cells that likely harbor the recruited bacteria. Alternatively, original microbiomes might not have been completely removed during antibiotic treatment, but this would require that remaining bacteria were sparse enough to evade detection with fluorescence microscopy and that they were unable to grow in marine broth tests. Overall, results demonstrate the *Phaeocystis* colonial microbiome is a consistent subset of co-cultured bacteria in media and is primarily composed of bacteria belonging to orders Alteromonadales, Burkholderiales, and Rhizobiales.

**Figure 4.**
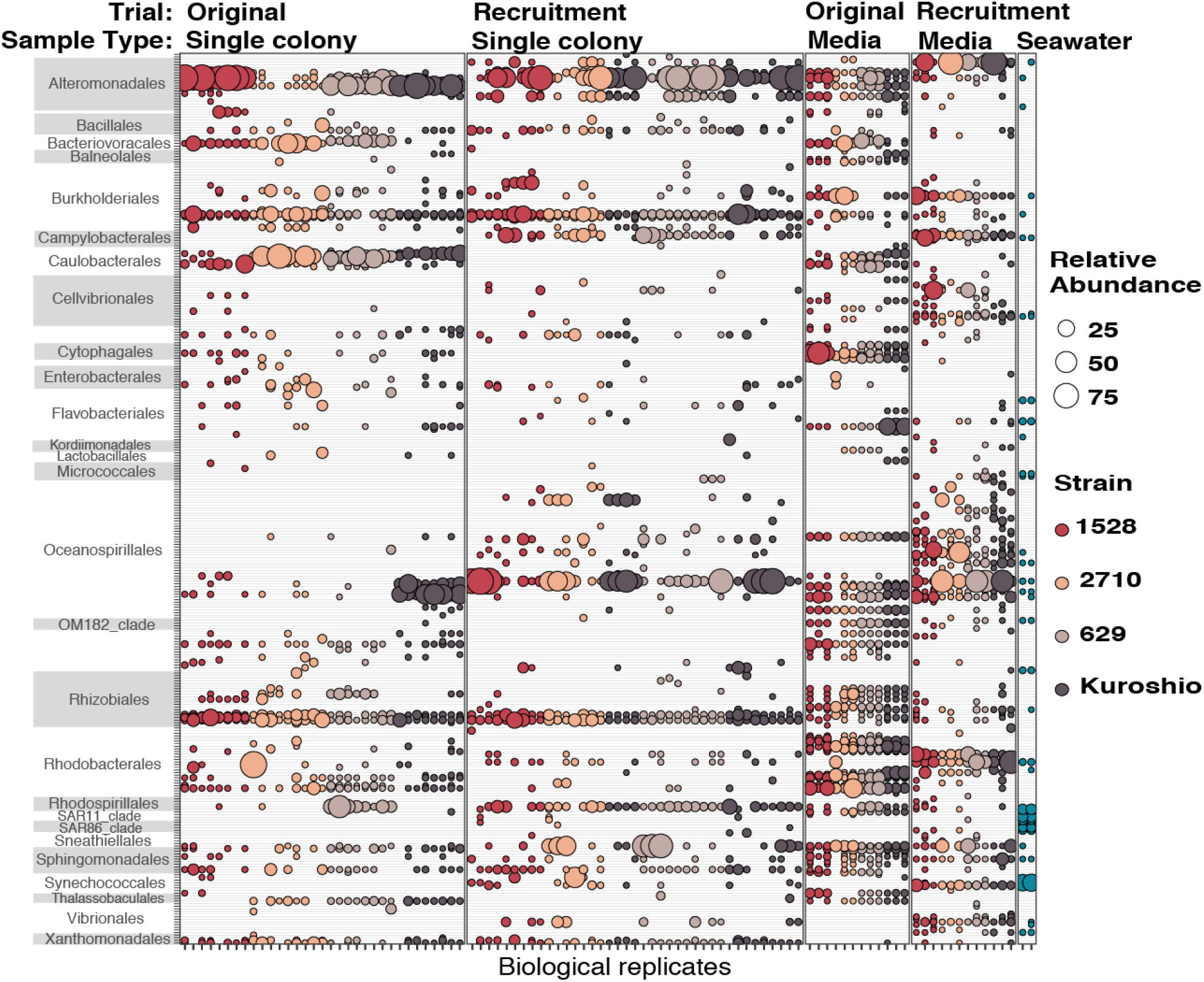
Relative abundance of Amplicon Sequence Variants (ASVs) detected in association with *Phaeocystis globosa* colonies and in culture media. Data were subset to include only ASVs that were present in at least one single-colony or media sample (*i.e*., not only present in seawater)—reducing the number of unique ASVs from 781 to 341—and the relative abundance of each remaining ASV in every sample is represented by bubble size. Bubbles are colored by *P. globosa* source strain or inoculating seawater and are grouped by trial (original or recruitment) and sample type (single colony, media, and inoculating seawater). Columns are biological replicates of each sample type and rows are individual ASVs. ASVs are grouped on the y-axis by taxonomic orders, which appear alphabetically.

### Impact of microbiomes on *Phaeocystis* growth when nitrate or B-vitamin deprived

Growth was monitored in axenic and nonaxenic culture replicates of each *P. globosa* strain in three media formulations: replete L1-Si with no modifications, L1-Si media with nitrate omitted, and L1-Si media with B-vitamin solution omitted (no added source of thiamine, biotin, or cobalamin). Growth was maintained in replete L1-Si media for all strains across the five transfers monitored (Fig. 5). In contrast, growth slowed for nonaxenic replicates of all four strains in L1-Si media without nitrate after the second transfer and ceased after the third transfer, but was unaffected in axenic replicates. Conversely, growth slowed for axenic replicates of all four strains in L1-Si media without B-vitamins after the fourth transfer, but was unaffected in nonaxenic replicates. These results suggest that when nitrogen is limiting, co-cultured bacteria compete for the limiting resource, whereas when vitamins become limiting, co-cultured bacteria mitigate limitation.

**Figure 5.**
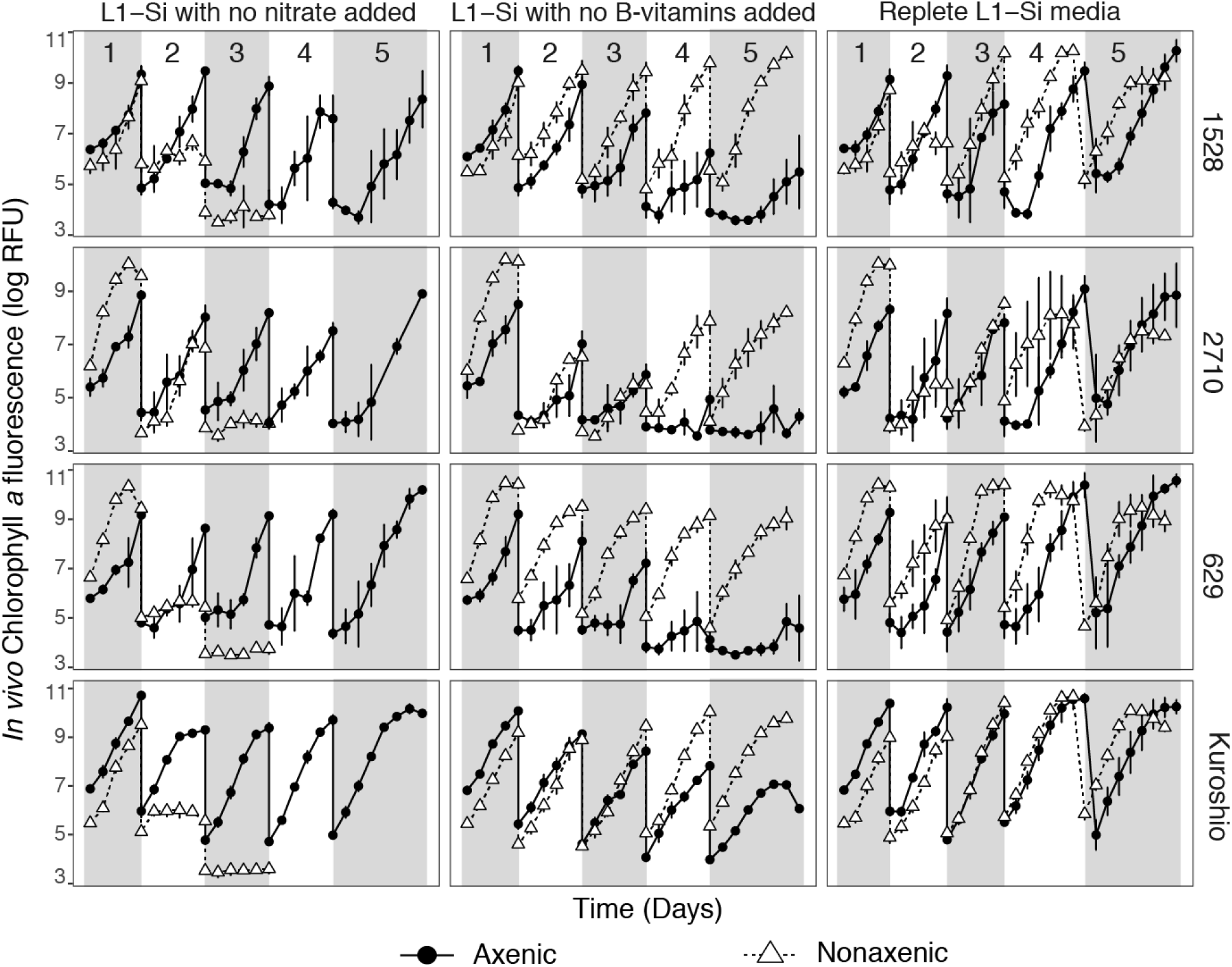
Growth curves for four strains of *Phaeocystis globosa* grown in L1-Si media with no nitrate added, L1-Si media with no B-vitamins added, and in replete L1-Si media. Each set of conditions were established in triplicate—each point is a mean of three biological replicates and error bars are one standard deviation of the mean—and cultures were maintained for five sequential transfers of five (transfers 1–4) to seven days (transfer 5). Serial transfers are denoted by alternating shaded regions on the plots and are numbered in the top panels. Growth was maintained in all replicates across all five transfers in replete L1-Si media; growth slowed after the 4th transfer in axenic replicates in L1-Si media without B-vitamins; growth slowed after the second transfer (and then ceased after the third) in nonaxenic replicates grown in L1-Si media without nitrate.

## Discussion

This study shows similarly composed bacterial communities are found in co-culture with *Phaeocystis globosa* strains that were isolated from different ocean basins across several decades (Fig. 1, 3, 4). Moreover, original communities directly associated with individual colonies were a predictable subset of co-cultured bacteria in culture media, and bacterial communities recruited to colonies converged towards the original composition of colony microbiomes (Fig. 4). The consistency of these communities is evidence *Phaeocystis globosa* colonies harbor a persistent core microbiome and selective processes drive microbiome assembly. Core constituents of original and recruited colony microbiomes included bacteria belonging to orders Alteromonadales, Burkholderiales, and Rhizobiales (Fig. 4). Of these, only Alteromonadales bacteria are core components of diatom phycosphere communities [20], but Rhizobiales bacteria have been found in association with the diatom *Asterionellopsis glacialis* [54]. Burkholderiales and Rhizobiales are commonly found in plant rhizospheres [55, 56]. Differences in *Phaeocystis* phycosphere community composition compared to other phytoplankton reflects the uniqueness of the *Phaeocystis* colony microenvironment. Furthermore, bacteria in co-culture with *P. globosa* mitigated nutrient stress when B-vitamins were withheld but accelerated culture collapse when nitrate was withheld (Fig. 5). This implies bacterial counterparts in the *Phaeocystis* phycosphere play a dual role alternating between mutualism and opportunism with their host [57].

### The *Phaeocystis globosa* microbiome is a stable-state system shaped by selective assembly mechanisms

Bacterial communities in co-culture with long-term *Phaeocystis* cultures are consistently composed and stable through time. The original media microbiomes were almost identical in *P. globosa* CCMP1528, CCMP2710, CCMP629, and Kuroshio strains, despite each of these strains being isolated from a different ocean basin during a different decade. Likewise, the original individual colony microbiomes were strikingly similar (Fig. 3, 4). However, despite sharing many core constituents (Fig. 4), overall compositions of original Kuroshio strain colony microbiomes were significantly different from the original colony microbiomes of the other strains (Fig. 3b). Since the Kuroshio strain was isolated most recently and was not sourced from a common culture bank, its different original microbiome suggests time and place of isolation influences the composition of *Phaeocystis* microbiomes—as reported for other phytoplankton [58]—but time spent in a common culture facility reduces initial variability. Convergence of colonial microbiomes of all four strains, including the Kuroshio strain, after antibiotic treatment and exposure to bacteria from seawater (Fig. 3b, 4) is further evidence that *Phaeocystis* microbiomes are stable systems shaped by selective, deterministic assembly mechanisms: given access to the same bacteria, all four strains recruit bacterial communities that are not significantly different from each other (*F* = 1.26, *R*^2^ = 0.09, *p* = 0.117).

Selection of *P. globosa* microbiome bacteria likely arises from the unique physical and chemical characteristics of colonies. First, the large size of colonies relative to individual *Phaeocystis* cells and other phytoplankton creates more potential habitat for bacteria and, therefore, increases the diversity that can be supported [59]. Second, the colony skin likely represents a physical barrier limiting access to the inner matrix [60], representing a first step in microbiome selection. Third, the colony matrix is made up of a unique medley of mucopolysaccharides, polysaccharides, and other organic molecules, including amino sugars [30, 31, 35], which would attract and support bacteria with unique enzymatic capabilities to exploit these resources [33]. Finally, additional community filtering may arise from feedback from and between microbiome bacteria, which can produce antibacterial compounds [61] or elicit phytoplankton defense responses [20]. For instance, the *Phaeocystis* colony matrix has high concentrations of acrylate, which has antimicrobial activity [27, 62]. Together, these *Phaeocystis* traits lead to consistent microbiome community assembly across individual colonies and across host strains, as was seen in this study. However, the convergence of recruited microbiomes and their differences from original microbiomes also demonstrate some functional redundancy in microbiome bacteria (*i.e*., phylogenetically distinct bacteria can occupy the same niche) [37].

### Core members of original and recruited *Phaeocystis globosa* colony microbiomes

Core members of the *Phaeocystis globosa* colony microbiome—bacteria with ASVs detected in almost all original and recruited colony microbiomes—belonged to orders Alteromonadales (family Alteromondacea), Burkholderiales (families Oxalobacteraceae and Comamonabacteraceae), and Rhizobiales (family Rhizobiaceae) (Fig. 4). Of these, only Alteromonadales bacteria are common inhabitants of diatom phycospheres [20]. In a BLAST search [63] against the NCBI nr/nt database (accessed: 10/4/2021, [64]), the three Alteromonadacea ASVs consistently abundant in colony microbiomes shared 99–100% nucleotide identity with sequences from *Alteromonas macleodii. A. macleodii—* a ubiquitous particle-associated marine copiotroph [65]—is an opportunistic, rather than symbiotic, member of diatom microbiomes [20] that directly competes with diatoms for nitrate [66]. Given how prevalent these bacteria are on marine particles and phytoplankton, it is unsurprising they are also associated with *Phaeocystis*. However, as with diatoms, we suspect the relationship between *Phaeocystis* and *A. macleodii* is more opportunistic than symbiotic in nature.

While bacteria in orders Burkholderiales and Rhizobiales are not commonly found living in association with other phytoplankton studied thus far, they are often associated with plant rhizospheres. The Burkholderiales ASV from the family Oxalobacteraceae that was consistently associated with *P. globosa* colonies shared 100% sequence identity with bacteria in the genus *Herbaspirillum*, which have been isolated from rhizospheres of many plants, and from soil and aquatic environments. *Herbaspirillum* strains have genomic potential to fix nitrogen, reduce nitrite, and produce siderophores [67]. *Herbaspirillum* strains also produce plant growth hormones—such as gibberellins and indole acetic acid [68]—and have been shown to promote rice, corn, and sugarcane growth [69]. The other core Burkholderiales ASV (family Comamonadaceae) shared 100% sequence identity with sequences deriving from the genus *Delftia. Delftia spp*. are considered Plant Growth-Promoting Rhizobacteria (PGPR), have capacity to produce siderophores [70], and can metabolize aromatic compounds, including benzene [71]. From the order Rhizobiales, one core ASV shared 100% identity with *Ochrobactrum pseudogrigonensis* sequences. *O. pseudogrigonensis* is a rhizosphere bacteria that promotes plant growth through induced stress tolerance, *e.g*., by producing antioxidants [56]. The other core Rhizobiales ASV shared 100% identity with *Rhizobium radiobacter*. Many *R. radiobacter* strains are plant pathogens, but nonpathogenic strains promote plant growth and can induce resistance to other pathogens [55]. While further work is needed to fully characterize how members of the core *Phaeocystis* microbiome influence *Phaeocystis* growth, our results indicate that, apart from *A. macleodii*, the core members of the *P. globosa* colony microbiome are broadly symbiotic bacteria with the potential to promote growth through phytohormones, nitrogen-cycling, or siderophore production.

### Impact of microbiomes on *Phaeocystis* growth under nitrogen and vitamin deprivation

Taken together, consistent and reproducible compositions of *P. globosa* microbiomes and the growth-promoting potential of most core microbiome members point toward beneficial interactions between *Phaeocystis* and microbiome bacteria. Interestingly, growth experiments involving axenic and nonaxenic *P. globosa* strains grown under nitrogen and vitamin deprivation revealed contrasting bacterial effects. Axenic cultures persisted longer than nonaxenic cultures when nitrate was withheld (Fig. 5), demonstrating competition between *Phaeocystis* and co-cultured bacteria for this important resource. This result aligns with the high relative abundance of *A. macleodii* in colony microbiomes (Fig. 4), as *A. macleodii* competes with phytoplankton for nitrate [66]. In a natural setting, these results might indicate microbiome interactions disadvantage *Phaeocystis* if nitrogen is limiting. However, given that *A. macleodii* is often also associated with diatoms [20], diatoms may be subject to similar ecological competition with heterotrophs. Indeed, the influence of nitrogen limitation on *Phaeocystis* and diatom bloom dynamics is not straightforward, and each group has come out on top when nitrogen was limiting in different ecosystems [23, 25, 72].

In contrast to when nitrate was withheld, axenic *P. globosa* growth was diminished compared to nonaxenic cultures when B-vitamins were withheld (Fig. 5), demonstrating co-cultured bacteria can produce essential B-vitamins (*i.e*., thiamine, cobalamin; [42]) and stave off growth-diminishing vitamin stress. This is significant because B-vitamins can be limiting in coastal areas where *Phaeocystis* blooms occur [41, 73, 74]. While several major heterotrophic cobalamin and thiamine synthesizing bacteria—namely Oceanospirillales [7], Rhodospirillales, and Flavobacteriales bacteria [75]—were in our dataset, they were more consistently present and abundant in media microbiomes than in colony microbiomes (Fig. 4). Nonetheless, two Oceanospirillales ASVs were abundant core components of original Kuroshio strain colonial microbiomes, and an Oceanospirillales and a Rhodospirillales ASV were both recruited at relatively high abundance to almost all colonies sampled in the recruitment trial (Fig. 4). The bacteria represented by these ASVs could indeed contribute to vitamin synthesis in *Phaeocystis* colony microbiomes. However, nutrient-limitation experiments only included *Phaeocystis* cultures retaining original microbiomes, where most recognized vitamin-producing bacteria were not tightly associated with colonies. Therefore, it is possible that other colony-associated bacteria are vitamin synthesizers or that vitamin synthesizers were more loosely associated with colonies (*i.e*., not attached to colony surfaces or inhabiting the colonial matrix). A wide diversity of marine bacteria are capable of vitamin synthesis [75, 76], thus necessitating evaluation of genomic potential and gene expression in *Phaeocystis* microbiomes to ultimately identify bacterial vitamin sources. Regardless of which microbiome constituents produce or consume limiting nutrients, challenging axenic and nonaxenic *P. globosa* with different limiting conditions clearly demonstrated interactions with the same total microbiome can be beneficial under one set of conditions and detrimental under another.

### Conclusions and Future Directions

Though interactions between phytoplankton and bacteria occur on small scales, they have large-scale repercussions for primary production and carbon cycling in aquatic ecosystems. A more complete understanding of these interactions that includes diverse phytoplankton taxa will improve our capabilities to model and predict changes to the marine environment. This study showed *Phaeocystis globosa* has consistent and reproducible core microbiomes co-cultured in media and associated with individual colonies. Members of the core microbiome have potential to produce growth-promoting phytohormones, fix nitrogen, synthesize vitamins, and release siderophores. Our experiments showed co-cultured bacteria provide vitamin resources and drawdown nitrate. Further work including metagenome and metatranscriptome sequencing to assess genomic potential and gene expression in *Phaeocystis* microbiomes is required to fully characterize mechanisms of interaction and currencies of exchange between *Phaeocystis* and bacteria. Additionally, future work should expand to other bloom-forming *Phaeocystis* species, determine how bacterial interactions impact *Phaeocystis* growth under other ecologically-relevant conditions—such as iron and phosphorus limitation—and investigate how increasing sea-surface temperatures associated with global climate change will impact microbiome interactions with *Phaeocystis* species.

## Acknowledgments

Research was funded by the Marine Biophysics Unit of the Okinawa Institute of Science and Technology (OIST) Graduate University. MMB was supported by an OIST Junior Research Fellowship, a Postdoctoral Scholarship granted by Woods Hole Oceanographic Institution, and a Simons Foundation Postdoctoral Fellowship in Marine Microbiology (award # 874439). We thank the captain and crew of the Okinawa Prefectural Fisheries and Ocean Research Center ship Tonan Maru for collecting seawater from the East China Sea for the culture of the Kuroshio strain of *Phaeocystis globosa*. We further thank Kazumi Inoha for coordinating seawater sampling on the Tonan Maru. Dawn Moran provided valuable advice for treating phytoplankton cultures with antibiotics. The OIST DNA sequencing section (Onna, Okinawa, Japan) performed the sequencing for this project.

## Supplemental Materials

**Figure S1.**
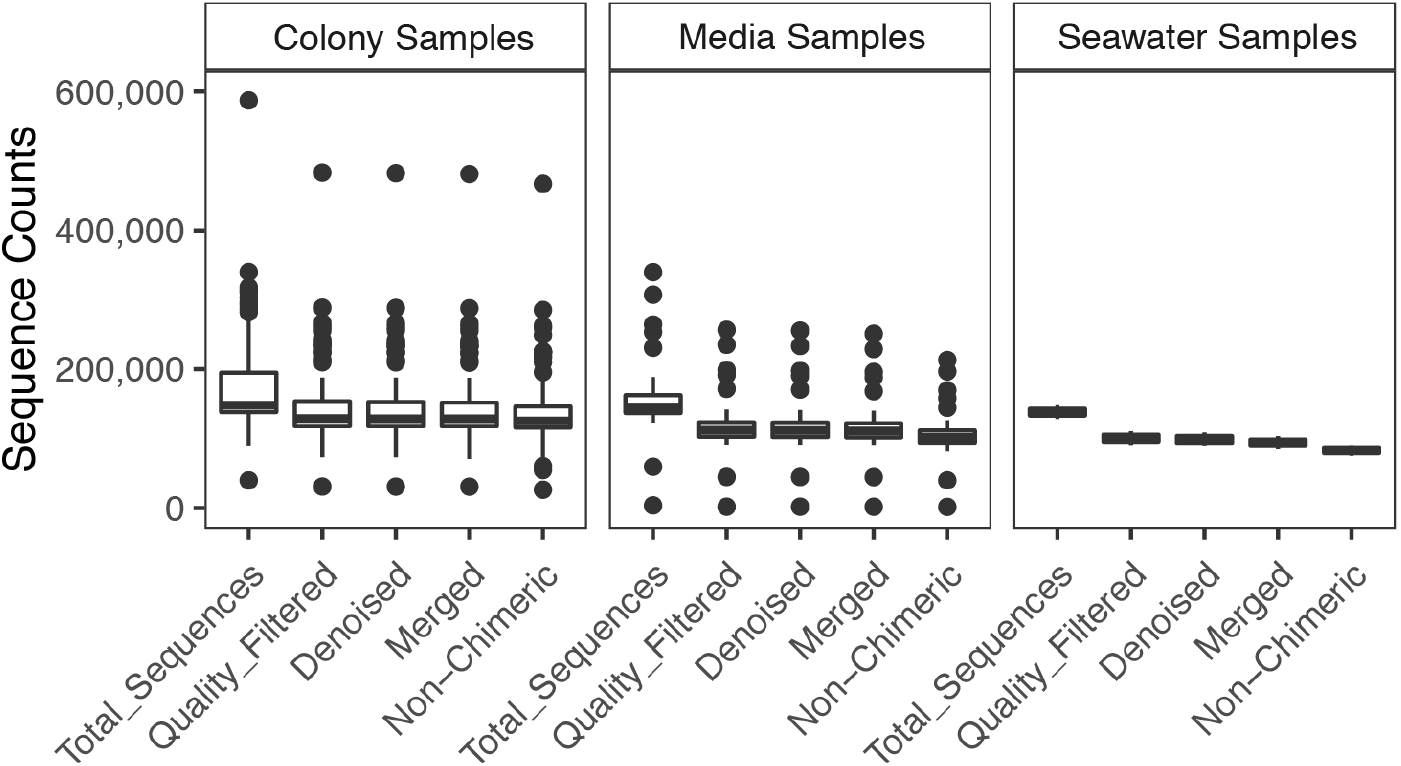
Total sequence read counts and number of remaining sequencing reads after each processing step in the DADA2 pipeline. The plot is faceted by sample type: single colony samples (*n* = 74), media samples (*n* = 24), and seawater samples (*n* = 2). Boxes denote the 25^th^ to 75^th^ percentile ranges for each sample type; the center bar is the median; whiskers show 1.5 times the interquartile range above the 75^th^ percentile and below the 25^th^ percentile; outlier points are values outside 1.5 times the interquartile range above the 75^th^ percentile and below the 25^th^ percentile. Raw sequence counts for each sample are available in table format through the following link: https://maggimars.github.io/phaeosphere/recruitmentStudy/kable.html.

**Figure S2.**
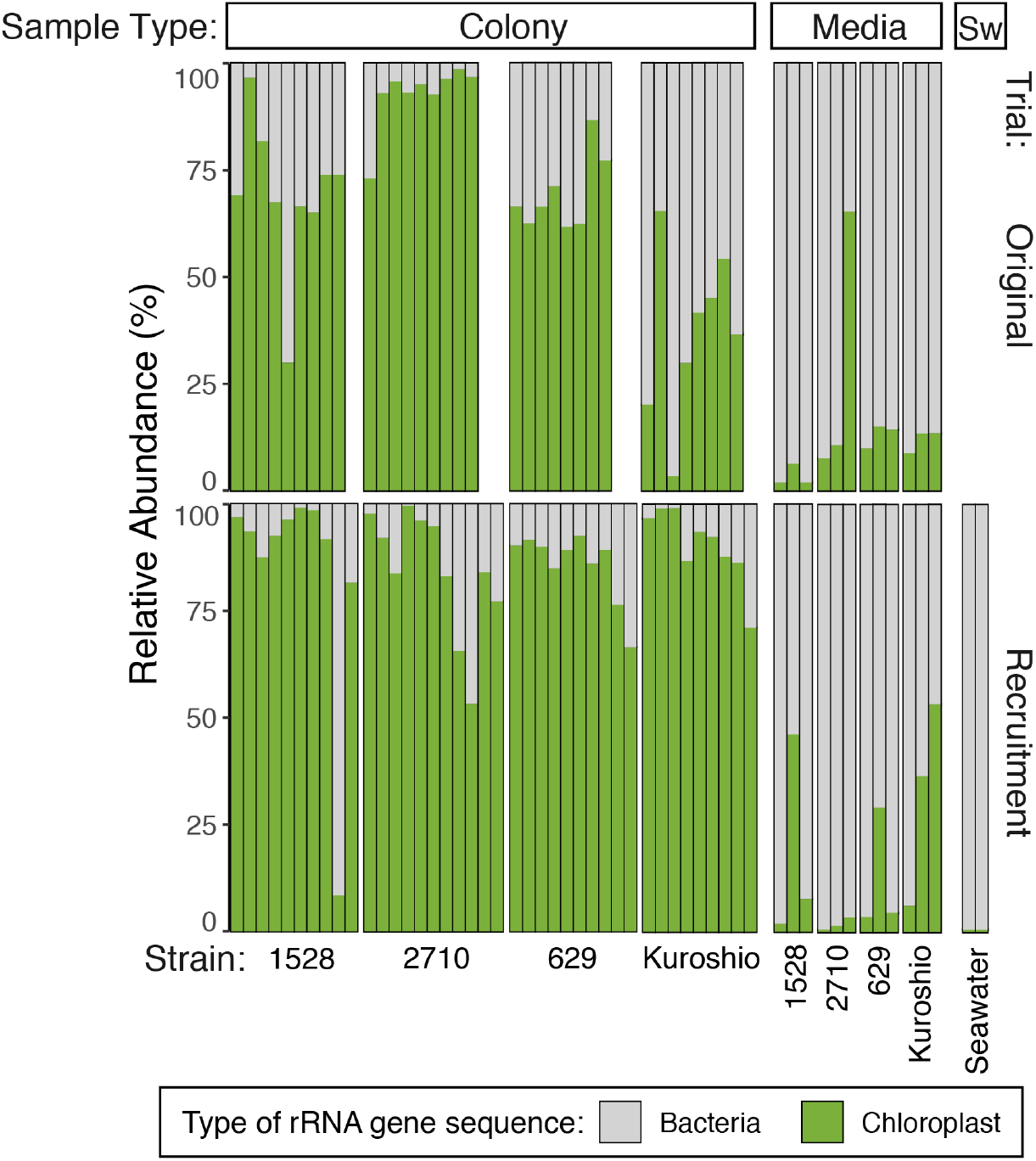
Relative abundance of bacterial and chloroplast 16S rRNA gene sequences in each sample. A mean of 22% of sequences or 31 095 sequences from colony samples were classified as bacteria. A mean of 85% of sequences or 93 042 sequences from media samples were classified as bacteria. The two sequencing replicates representing the natural bacterial assemblage at the start of the recruitment trial had the largest mean proportion of sequencing reads classified as bacteria: 99.5% (mean 82 219 sequences per sample).

**Figure S3.**
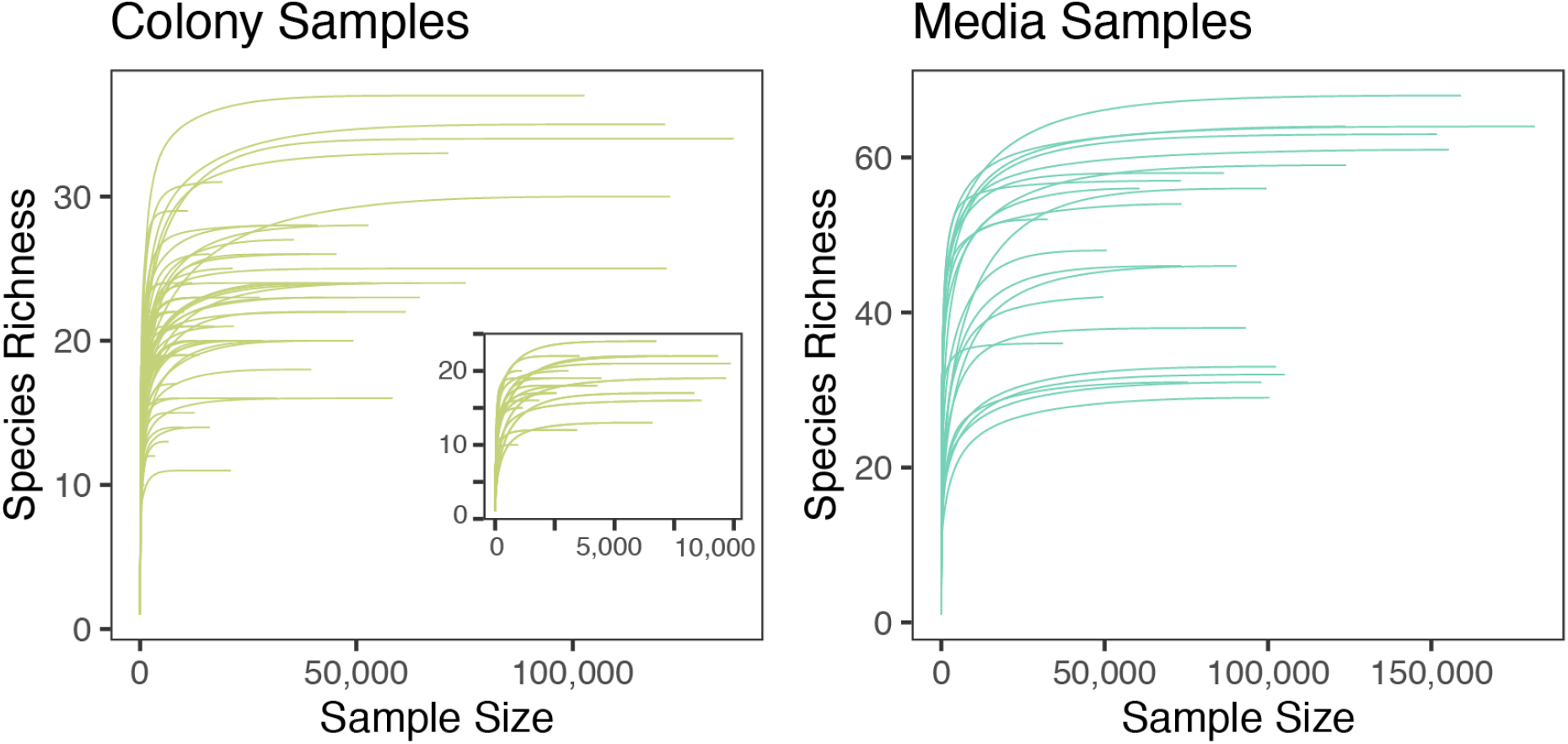
Rarefaction curves for single-colony and media microbiome samples. Rarefaction was performed with phyloseq after chloroplast sequences were removed from the data and results were plotted with the function ‘ggrare’. The colony sample inset plot shows samples with less than 10 000 sequences remaining after chloroplast sequences were removed. All samples reach species (ASV) richness saturation.

## References

1. Moran MA, Kujawinski EB, Stubbins A, Fatland R, Aluwihare LI, Buchan A, et al. Deciphering ocean carbon in a changing world. Proc Natl Acad Sci USA 2016; 113:3143–3151.

2. Seymour JR, Amin SA, Raina J-B, Stocker R. Zooming in on the phycosphere: the ecological interface for phytoplankton–bacteria relationships. Nature Microbiology 2017; 2

3. Mendes R, Kruijt M, de Bruijn I, Dekkers E, van der Voort M, Schneider JHM, et al. Deciphering the rhizosphere microbiome for disease-suppressive bacteria. Science 2011;332: 1097–1100.

4. Cirri E, Pohnert G. Algae-bacteria interactions that balance the planktonic microbiome. New Phytol 2019; 223: 100–106.

5. Amin SA, Hmelo LR, van Tol HM, Durham BP, Carlson LT, Heal KR, et al. Interaction and signaling between a cosmopolitan phytoplankton and associated bacteria. Nature 2015;522: 98–101.

6. Grant MAA, Kazamia E, Cicuta P, Smith AG. Direct exchange of vitamin B_12_ is demonstrated by modelling the growth dynamics of algal-bacterial cocultures. ISME J 2014;8: 1418–1427.

7. Bertrand EM, McCrow JP, Moustafa A, Zheng H, McQuaid JB, Delmont TO, et al. Phytoplankton–bacterial interactions mediate micronutrient colimitation at the coastal Antarctic sea ice edge. Proc Natl Acad Sci USA 2015; 112: 9938–9943

8. Durham BP, Sharma S, Luo H, Smith CB, Amin SA, Bender SJ, et al. Cryptic carbon and sulfur cycling between surface ocean plankton. Proc Natl Acad Sci USA 2015; 112:453–457.

9. Suleiman M, Zecher K, Yücel O, Jagmann N, Philipp B. Interkingdom Cross-Feeding of Ammonium from Marine Methylamine-Degrading Bacteria to the Diatom Phaeodactylum tricornutum. Appl Environ Microbiol 2016; 82: 7113–7122.

10. Seyedsayamdost MR, Case RJ, Kolter R, Clardy J. The Jekyll-and-Hyde chemistry of Phaeobacter gallaeciensis. Nature Chemistry 2011; 3: 331–335

11. Ratnarajah L, Blain S, Boyd PW, Fourquez M, Obernosterer I, Tagliabue A. Resource Colimitation Drives Competition Between Phytoplankton and Bacteria in the Southern Ocean. Geophys Res Lett 2021; 48: e2020GL088369.

12. Løvdal T, Eichner C, Grossart H-P, Carbonnel V, Chou L, Martin-Jézéquel V, et al. Competition for inorganic and organic forms of nitrogen and phosphorous between phytoplankton and bacteria during an Emiliania huxleyi spring bloom. Biogeosciences 2008;5: 371–383.

13. Arrigo KR, Robinson DH, Worthen DL, Dunbar RB, DiTullio GR, VanWoert M, et al. Phytoplankton community structure and the drawdown of nutrients and CO_2_ in the southern ocean. Science 1999; 283: 365–367.

14. Geider R, La Roche J. Redfield revisited: variability of C:N:P in marine microalgae and its biochemical basis. European Journal of Phycology 2002; 37: 1–17

15. Smayda TJ. Normal and accelerated sinking of phytoplankton in the sea. Marine Geology 1971;11: 105–122

16. Amin SA, Parker MS, Armbrust EV. Interactions between diatoms and bacteria. Microbiol Mol Biol Rev 2012; 76: 667–684.

17. Tréguer P, Bowler C, Moriceau B, Dutkiewicz S, Gehlen M, Aumont O, et al. Influence of diatom diversity on the ocean biological carbon pump. Nature Geoscience 2018; 11: 27–37

18. Ferrer-González FX, Widner B, Holderman NR, Glushka J, Edison AS, Kujawinski EB, et al. Resource partitioning of phytoplankton metabolites that support bacterial heterotrophy. ISME J 2021; 15: 762–773.

19. Mönnich J, Tebben J, Bergemann J, Case R, Wohlrab S, Harder T. Niche-based assembly of bacterial consortia on the diatom Thalassiosira rotula is stable and reproducible. ISME J 2020; 14: 1614–1625.

20. Shibl AA, Isaac A, Ochsenkühn MA, Cárdenas A, Fei C, Behringer G, et al. Diatom modulation of select bacteria through use of two unique secondary metabolites. Proc Natl Acad Sci USA 2020; 117: 27445–27455.

21. Schoemann V, Becquevort S, Stefels J, Rousseau V, Lancelot C. Phaeocystis blooms in the global ocean and their controlling mechanisms: a review. J Sea Res 2005; 53: 43–66.

22. Peperzak L, Colijn F, Gieskes WWC, Peeters JCH. Development of the diatom-Phaeocystis spring bloom in the Dutch coastal zone of the North Sea: the silicon depletion versus the daily irradiance threshold hypothesis. J Plankton Res 1998; 20: 517–537.

23. Hai D-N, Lam N-N, Dippner JW. Development of Phaeocystis globosa blooms in the upwelling waters of the South Central coast of Viet Nam. J Mar Syst 2010; 83: 253–261.

24. Wang X, Song H, Wang Y, Chen N. Research on the biology and ecology of the harmful algal bloom species Phaeocystis globosa in China: Progresses in the last 20 years. Harmful Algae 2021; 107: 102057.

25. Jiang M, Borkman DG, Scott Libby P, Townsend DW, Zhou M. Nutrient input and the competition between Phaeocystis pouchetii and diatoms in Massachusetts Bay spring bloom. J Mar Syst 2014; 134: 29–44.

26. Nissen C, Vogt M. Factors controlling the competition between Phaeocystis and diatoms in the Southern Ocean and implications for carbon export fluxes. Biogeosciences 2021; 18:251–283.

27. Mars Brisbin M, Mitarai S. Differential Gene Expression Supports a Resource-Intensive,Defensive Role for Colony Production in the Bloom-Forming Haptophyte, Phaeocystis globosa. J Eukaryot Microbiol 2019; 66: 788–801.

28. Zhu Z, Meng R, Smith WO Jr, Doan-Nhu H, Nguyen-Ngoc L, Jiang X. Bacterial Composition Associated With Giant Colonies of the Harmful Algal Species Phaeocystis globosa. Front Microbiol 2021; 12: 737484.

29. Delmont TO, Hammar KM, Ducklow HW, Yager PL, Post AF. Phaeocystis antarctica blooms strongly influence bacterial community structures in the Amundsen Sea polynya. Front Microbiol 2014; 5: 646.

30. Verity PG, Whipple SJ, Nejstgaard JC, Alderkamp A-C. Colony size, cell number, carbon and nitrogen contents of Phaeocystis pouchetii from western Norway. J Plankton Res 2007;29: 359–367.

31. Alderkamp A-C, Buma AGJ, van Rijssel M. The carbohydrates of Phaeocystis and their degradation in the microbial food web. Biogeochemistry 2007; 83: 99–118.

32. Smriga S, Fernandez VI, Mitchell JG, Stocker R. Chemotaxis toward phytoplankton drives organic matter partitioning among marine bacteria. Proc Natl Acad Sci USA 2016; 113:1576–1581.

33. Mühlenbruch M, Grossart H-P, Eigemann F, Voss M. Mini-review: Phytoplankton-derived polysaccharides in the marine environment and their interactions with heterotrophic bacteria. Environ Microbiol 2018; 20: 2671–2685.

34. Raina J-B, Fernandez V, Lambert B, Stocker R, Seymour JR. The role of microbial motility and chemotaxis in symbiosis. Nat Rev Microbiol 2019; 17: 284–294.

35. Solomon CM, Lessard EJ, Keil RG, Foy MS. Characterization of extracellular polymers of Phaeocystis globosa and P. antarctica. Mar Ecol Prog Ser 2003; 250: 81–89.

36. Shen P, Qi Y, Wang Y, Huang L. Phaeocystis globosa Scherffel, a harmful microalga, and its production of dimethylsulfoniopropionate. Chin J Oceanol Limnol 2011; 29: 869–873.

37. Louca S, Polz MF, Mazel F, Albright MBN, Huber JA, O’Connor MI, et al. Function and functional redundancy in microbial systems. Nat Ecol Evol 2018; 2: 936–943.

38. Wang J, Bouwman AF, Liu X, Beusen AHW, Van Dingenen R, Dentener F, et al. Harmful Algal Blooms in Chinese Coastal Waters Will Persist Due to Perturbed Nutrient Ratios. Environ Sci Technol Lett 2021; 8: 276–284.

39. Foster RA, Kuypers MMM, Vagner T, Paerl RW, Musat N, Zehr JP. Nitrogen fixation and transfer in open ocean diatom-cyanobacterial symbioses. ISME J 2011; 5: 1484–1493.

40. Helliwell KE. The roles of B vitamins in phytoplankton nutrition: new perspectives and prospects. New Phytol 2017; 216: 62–68.

41. Bertrand EM, Saito MA, Rose JM, Riesselman CR, Lohan MC, Noble AE, et al. Vitamin B_12_ and iron colimitation of phytoplankton growth in the Ross Sea. Limnol Oceanogr 2007; 52:1079–1093.

42. Tang YZ, Koch F, Gobler CJ. Most harmful algal bloom species are vitamin B_1_ and B_12_ auxotrophs. Proc Natl Acad Sci USA 2010; 107: 20756–20761.

43. Croft MT, Lawrence AD, Raux-Deery E, Warren MJ, Smith AG. Algae acquire vitamin B_12_ through a symbiotic relationship with bacteria. Nature 2005; 438: 90–93.

44. Guillard RRL, Hargraves PE. Stichochrysis immobilis is a diatom, not a chrysophyte. Phycologia 1993; 32: 234–236

45. Hamilton PB, Lefebvre KE, Bull RD. Single cell PCR amplification of diatoms using fresh and preserved samples. Front Microbiol 2015; 6: 1084.

46. Callahan BJ, McMurdie PJ, Rosen MJ, Han AW, Johnson AJA, Holmes SP. DADA2:High-resolution sample inference from Illumina amplicon data. Nat Methods 2016; 13:581–583.

47. Bolyen E, Rideout JR, Dillon MR, Bokulich NA, Abnet CC, Al-Ghalith GA, et al. Reproducible, interactive, scalable and extensible microbiome data science using QIIME 2. Nat Biotechnol 2019; 37: 852–857.

48. Quast C, Pruesse E, Yilmaz P, Gerken J, Schweer T, Glo FO, et al. The SILVA ribosomal RNA gene database project: improved data processing and web-based tools. 2013; 41:590–596.

49. Bokulich NA, Kaehler BD, Rideout JR, Dillon M, Bolyen E, Knight R, et al. Optimizing taxonomic classification of marker-gene amplicon sequences with QIIME 2’s q2-feature-classifier plugin. Microbiome 2018; 6: 90.

50. R Core Team. R: A language and environment for statistical computing. 2018.

51. Mcmurdie PJ, Holmes S. phyloseq : An R Package for Reproducible Interactive Analysis and Graphics of Microbiome Census Data. 2013; 8.

52. Gloor GB, Macklaim JM, Pawlowsky-Glahn V, Egozcue JJ. Microbiome Datasets Are Compositional: And This Is Not Optional. Front Microbiol 2017; 8: 2224.

53. Oksanen J, Guillaume Blanchet F, Friendly M, Kindt R, Legendre P, McGlinn D, et al. vegan: Community Ecology Package. R package version 2.5-4. 2019.

54. Behringer G, Ochsenkühn MA, Fei C, Fanning J, Koester JA, Amin SA. Bacterial Communities of Diatoms Display Strong Conservation Across Strains and Time. Front Microbiol 2018; 9: 659.

55. Glaeser SP, Imani J, Alabid I, Guo H, Kumar N, Kämpfer P, et al. Non-pathogenic Rhizobium radiobacter F4 deploys plant beneficial activity independent of its host Piriformospora indica. ISME J 2016; 10: 871–884.

56. Chakraborty U, Chakraborty BN, Dey PL, Chakraborty AP, Sarkar J. Biochemical responses of wheat plants primed with Ochrobactrum pseudogrignonense and subjected to salinity stress. Agric Res 2019; 8: 427–440.

57. Johnson WM, Alexander H, Bier RL, Miller DR, Muscarella ME, Pitz KJ, et al. Auxotrophic interactions: a stabilizing attribute of aquatic microbial communities? FEMS Microbiol Ecol 2020; 96.

58. Ajani PA, Kahlke T, Siboni N, Carney R, Murray SA, Seymour JR. The Microbiome of the Cosmopolitan Diatom Leptocylindrus Reveals Significant Spatial and Temporal Variability. Front Microbiol 2018; 9: 2758.

59. Connor EF, McCoy ED. The Statistics and Biology of the Species-Area Relationship. Am Nat 1979; 113: 791–833.

60. Hamm CE, Simson DA, Merkel R, Smetacek V. Colonies of Phaeocystis globosa are protected by a thin but tough skin. Mar Ecol Prog Ser 1999; 187: 101–111.

61. Geddes BA, Paramasivan P, Joffrin A, Thompson AL, Christensen K, Jorrin B, et al. Engineering transkingdom signalling in plants to control gene expression in rhizosphere bacteria. Nat Commun 2019; 10: 3430.

62. Sieburth JM. Acrylic acid, an‘ antibiotic’ principle in Phaeocystis blooms in Antarctic waters. Science 1960; 132: 676–677.

63. Camacho C, Coulouris G, Avagyan V, Ma N, Papadopoulos J, Bealer K, et al. BLAST+:architecture and applications. BMC Bioinformatics 2009; 10: 421.

64. Clark K, Karsch-Mizrachi I, Lipman DJ, Ostell J, Sayers EW. GenBank. Nucleic Acids Res 2016; 44: D67–72.

65. López-Pérez M, Gonzaga A, Martin-Cuadrado A-B, Onyshchenko O, Ghavidel A, Ghai R, et al. Genomes of surface isolates of Alteromonas macleodii: the life of a widespread marine opportunistic copiotroph. Sci Rep 2012; 2: 696.

66. Diner RE, Schwenck SM, McCrow JP, Zheng H, Allen AE. Genetic Manipulation of Competition for Nitrate between Heterotrophic Bacteria and Diatoms. Front Microbiol 2016;7: 880.

67. Monteiro RA, Balsanelli E, Wassem R, Marin AM, Brusamarello-Santos LCC, Schmidt MA, et al. Herbaspirillum-plant interactions: microscopical, histological and molecular aspects. Plant Soil 2012; 356: 175–196.

68. Bastián F, Cohen A, Piccoli P, Luna V, Baraldi R. Production of indole-3-acetic acid and gibberellins A1 and A3 by Acetobacter diazotrophicus and Herbaspirillum seropedicae in chemically-defined culture media. Plant Growth Regul 1998.

69. Gyaneshwar P, James EK, Reddy PM. Herbaspirillum colonization increases growth and nitrogen accumulation in aluminium-tolerant rice varieties. New Phytol 2002; 154: 131–145.

70. Guo H, Yang Y, Liu K, Xu W, Gao J, Duan H, et al. Comparative Genomic Analysis of Delftia tsuruhatensis MTQ3 and the Identification of Functional NRPS Genes for Siderophore Production. Biomed Res Int 2016; 2016: 3687619.

71. Vásquez-Piñeros MA, Martínez-Lavanchy PM, Jehmlich N, Pieper DH, Rincón CA, Harms H, et al. Delftia sp. LCW, a strain isolated from a constructed wetland shows novel properties for dimethylphenol isomers degradation. BMC Microbiol 2018; 18: 108.

72. Riegman R, Noordeloos AAM, Cadée GC. Phaeocystis blooms and eutrophication of the continental coastal zones of the North Sea. Mar Biol 1992; 112: 479–484.

73. Sañudo-Wilhelmy SA, Cutter LS, Durazo R, Smail EA, Gómez-Consarnau L, Webb EA, et al. Multiple B-vitamin depletion in large areas of the coastal ocean. Proc Natl Acad Sci USA 2012; 109: 14041–14045.

74. Gobler CJ, Norman C, Panzeca C, Taylor GT, Sañudo-Wilhelmy SA. Effect of B-vitamins (B_1_, B_12_) and inorganic nutrients on algal bloom dynamics in a coastal ecosystem. Aquat Microb Ecol 2007; 49: 181–194.

75. Gómez-Consarnau L, Sachdeva R, Gifford SM, Cutter LS, Fuhrman JA, Sañudo-Wilhelmy SA, et al. Mosaic patterns of B-vitamin synthesis and utilization in a natural marine microbial community. Environ Microbiol 2018; 20: 2809–2823.

76. Bertrand EM, Saito MA, Jeon YJ, Neilan BA. Vitamin B_12_ biosynthesis gene diversity in the Ross Sea: the identification of a new group of putative polar B_12_ biosynthesizers. Environ Microbiol 2011; 13: 1285–1298.

